# *nf-core*: Community curated bioinformatics pipelines

**DOI:** 10.1101/610741

**Authors:** Philip A Ewels, Alexander Peltzer, Sven Fillinger, Johannes Alneberg, Harshil Patel, Andreas Wilm, Maxime Ulysse Garcia, Paolo Di Tommaso, Sven Nahnsen

## Abstract

The standardization, portability, and reproducibility of analysis pipelines is a renowned problem within the bioinformatics community. Most pipelines are designed for execution on-premise, and the associated software dependencies are tightly coupled with the local compute environment. This leads to poor pipeline portability and reproducibility of the ensuing results - both of which are fundamental requirements for the validation of scientific findings. Here, we introduce *nf-core*: a framework that provides a community-driven, peer-reviewed platform for the development of best practice analysis pipelines written in Nextflow. Key obstacles in pipeline development such as portability, reproducibility, scalability and unified parallelism are inherently addressed by all *nf-core* pipelines. We are also continually developing a suite of tools that assist in the creation and development of both new and existing pipelines. Our primary goal is to provide a platform for high-quality, reproducible bioinformatics pipelines that can be utilized across various institutions and research facilities.

---

Being able to reproduce scientific results is the central tenet of the scientific method. However, moving towards FAIR (Findable, Accessible, Interoperable and Reusable) research methods^1^ in data-driven science is complex^2,3^. Central repositories such as qPortal^4^, omictools^5^, and the Galaxy toolshed^6^ make it possible to find existing pipelines and their associated tools. However, it is still notoriously challenging to develop analysis pipelines that are fully reproducible and interoperable across multiple systems and institutions - primarily due to differences in hardware, operating systems and software versions.

Although the recommended guidelines for some analysis pipelines have become standardised (e.g. GATK best practices^7^), the actual implementations are usually developed by individual researchers. As such, they are rarely thoroughly tested, well documented or written in a format whereby they can be shared with others. This not only hampers sustainable sharing of data and tools but also results in a proliferation of heterogeneous analysis pipelines, making it difficult for newcomers to find what they need to address a specific analysis question.

As the scale of ‘omics data and its associated analytical tools has grown dramatically, the scientific community is increasingly moving towards the use of specialized workflow management systems to build analysis pipelines. They separate the requirements of the underlying computing infrastructure from the analysis and workflow description, and therefore introduce a higher degree of portability compared to custom in-house scripts. One such popular tool is Nextflow^8^. Using Nextflow, software packages can be bundled with analysis pipelines using built-in integration of package managers like Conda, and container environments such as Docker and Singularity. Moreover, support for most common High Performance Computing (HPC) batch schedulers and cloud providers allows simple deployment of analysis pipelines on almost any infrastructure in a transparent and portable manner.

Other high-level approaches to facilitate the creation of analysis pipelines are available: Flowcraft^9^ and Pipeliner^10^ are meta-tools to dynamically build Nextflow pipelines from template blocks. The awesome-pipelines website^11^ provides a curated list of Nextflow pipelines developed by the community, however, these are highly individualised, and do not follow consistent implementation patterns. Other workflow tools have collections of pipelines, although with less extensively defined rules than *nf-core*. Snakemake Workflows^12^ is an early-stage community project attempting to collect best practice analysis pipelines based on the Snakemake^13^ workflow manager, the Galaxy Toolshed^6^ collects tools and workflows for use with Galaxy^14^, and the ENCODE project pipelines^15^ have established best-practice scripts and workflows for a number of genomics assay-types.

*nf-core* pipelines are built and maintained by an entire user community of researchers that all contribute to a larger objective, rather than a single group or Institute. While being strongly represented in the analysis of genomics data, the community also provides a more general framework for building and maintaining analysis pipelines in proteomics, metabolomics, and other life science applications. In theory, the same should also be applicable to the broader field of scientific data analysis^17–19^ including more data-intensive fields such as chemistry and physics.

Pipelines within *nf-core* all adhere to the same principles. These are described on the project homepage and cover topics such as software dependencies, code structure and the use of minimal datasets for testing purposes. Once a researcher is familiar with one *nf-core* analysis pipeline, all others will work on the same system with a similar logic. Detailed documentation and templates for the addition of new pipelines make it trivial to start a new project, making *nf-core* an excellent entry point for Nextflow pipeline development.

The driving force behind *nf-core* is its community. In addition to a curated collection of analysis pipelines, the *nf-core* community strives to provide a framework for developer best-practices, code standardization, and comprehensive documentation. The open nature of this community allows discussion and collaboration on both general best-practices and domain-specific pipeline implementation from experts across the world. Technical issues are typically resolved and reviewed by a group of people, ensuring that the resulting solutions are reliable, well designed and thoroughly tested. This open process breaks down traditional barriers and silos between research groups, resulting in high-quality pipelines that anyone can use. The early feedback and successive reviewing process allows developers to improve their analysis pipelines and ensures that they adhere to community standards. Through discussions and collaborative efforts, new guidelines and tools naturally evolve and are made available to all *nf-core* developers.

The hub of the *nf-core* community is its website (https://nf-co.re). It contains a list of all analysis pipelines as well as documentation, tutorials and communication platforms serving as the first point of entry for newcomers, and a point of reference for more experienced developers. All code is hosted on GitHub in the *nf-core* organization (https://github.com/nf-core/).

The *nf-core* project ties in well with other communities, notably Nextflow^8^, Bioconda^20^ and conda-forge^21^. Wherever possible, we encourage users and developers to contribute to these projects, such as packaging missing software in the bioconda or conda-forge repositories, helping the wider bioinformatics community.

Currently, *nf-core* encompasses a total of 27 analysis pipelines that span multiple life-science domains and analysis types. For example; `nf-core/rnaseq`^22^ generates gene counts and quality metrics for RNA-seq data, `nf-core/mhcquant`^23^ identifies and quantifies peptides from mass spectrometry raw data, `nf-core/sarek`^24^ can detect germline or somatic variants from normal or tumour/normal whole-genome or targeted sequencing data, and `nf-core/eager`^25,26^ processes genomics data from ancient DNA samples. On release, *nf-core* pipelines are assigned a unique version tag that links the pipeline code to its associated software dependencies. This provides users with the reassurance that the results will be fully reproducibility.

All *nf-core* pipelines are released under the highly permissive MIT license, encouraging downstream modification and reuse. Indeed, a key feature of *nf-core* analysis pipelines is to provide “base” pipelines that can be built upon for more niche project-specific outputs. All pipelines are required to provide high-quality documentation: both pipeline usage documentation with examples, and also a description of the generated results files.

Since *nf-core* pipelines use Nextflow, they can be deployed across a huge range of environments. All of the software dependencies required for any given analysis pipeline are available using either Conda, Docker or Singularity. A centralized repository holding Institute-specific configuration files means that a custom cluster-specific configuration profile can be made available for all pipelines instantly, through a single update.

**Figure 1:**
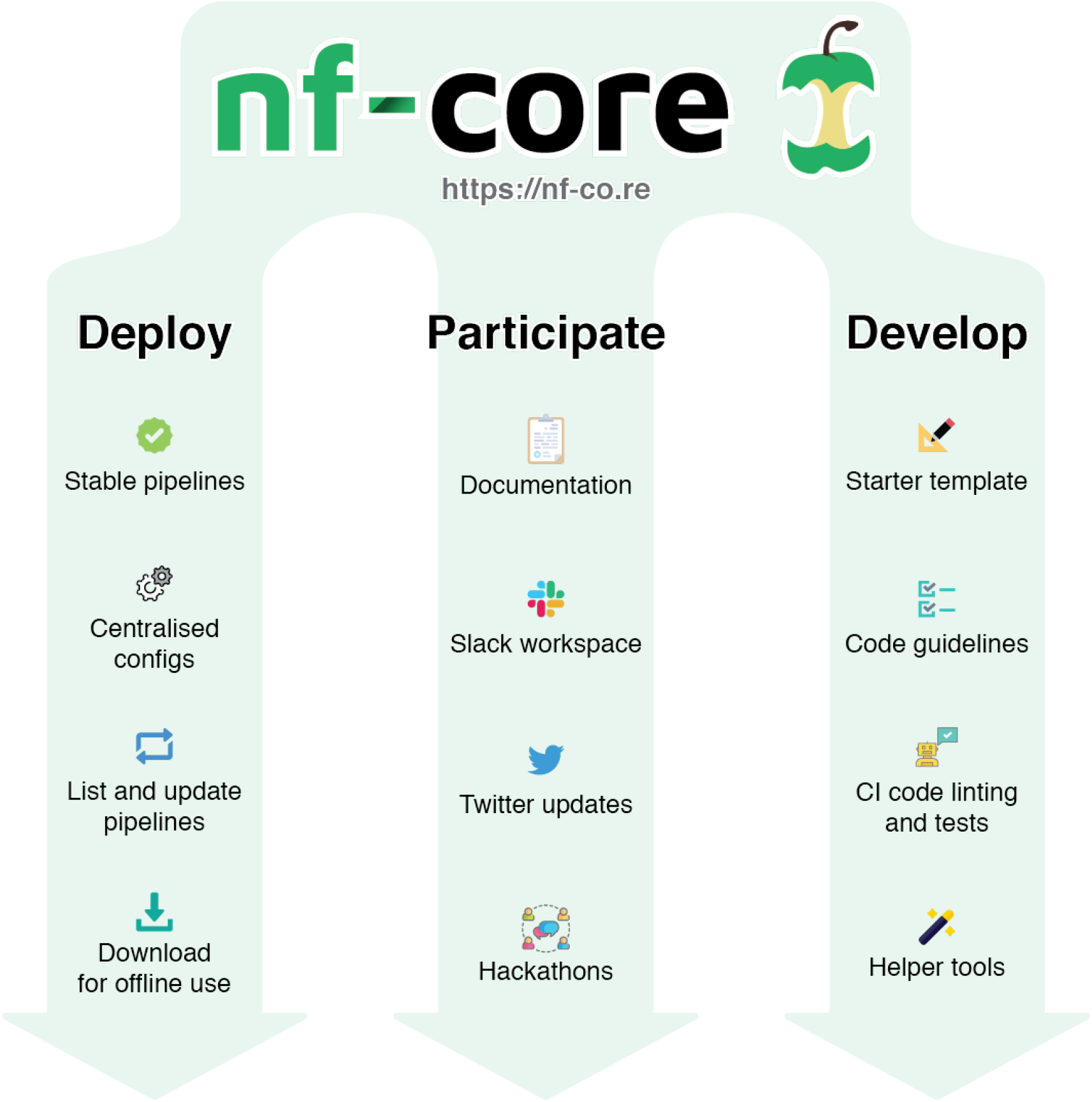
Main concepts of nf-core

To help *nf-core* users and developers, we provide a set of command-line tools to automate common tasks. Users can list available analysis pipelines and check that their versions are up to date (Fig. 2), as well as downloading all required files for offline use in a single command. Developers have a suite of commands to build, test and release new analysis pipelines, including a conformance check that validates code against current best-practice recommendations defined by the *nf-core* community (Table 1). Additional checks also provide helpful warnings, such as notifications about new versions of software available on conda-forge or bioconda.

Framework integration tests and test-dataset runs are automatically triggered whenever there is a change to a pipeline, using continuous integration with Travis CI^27^. This ensures that developers can guarantee a working pipeline at all times.

**Figure 2:**
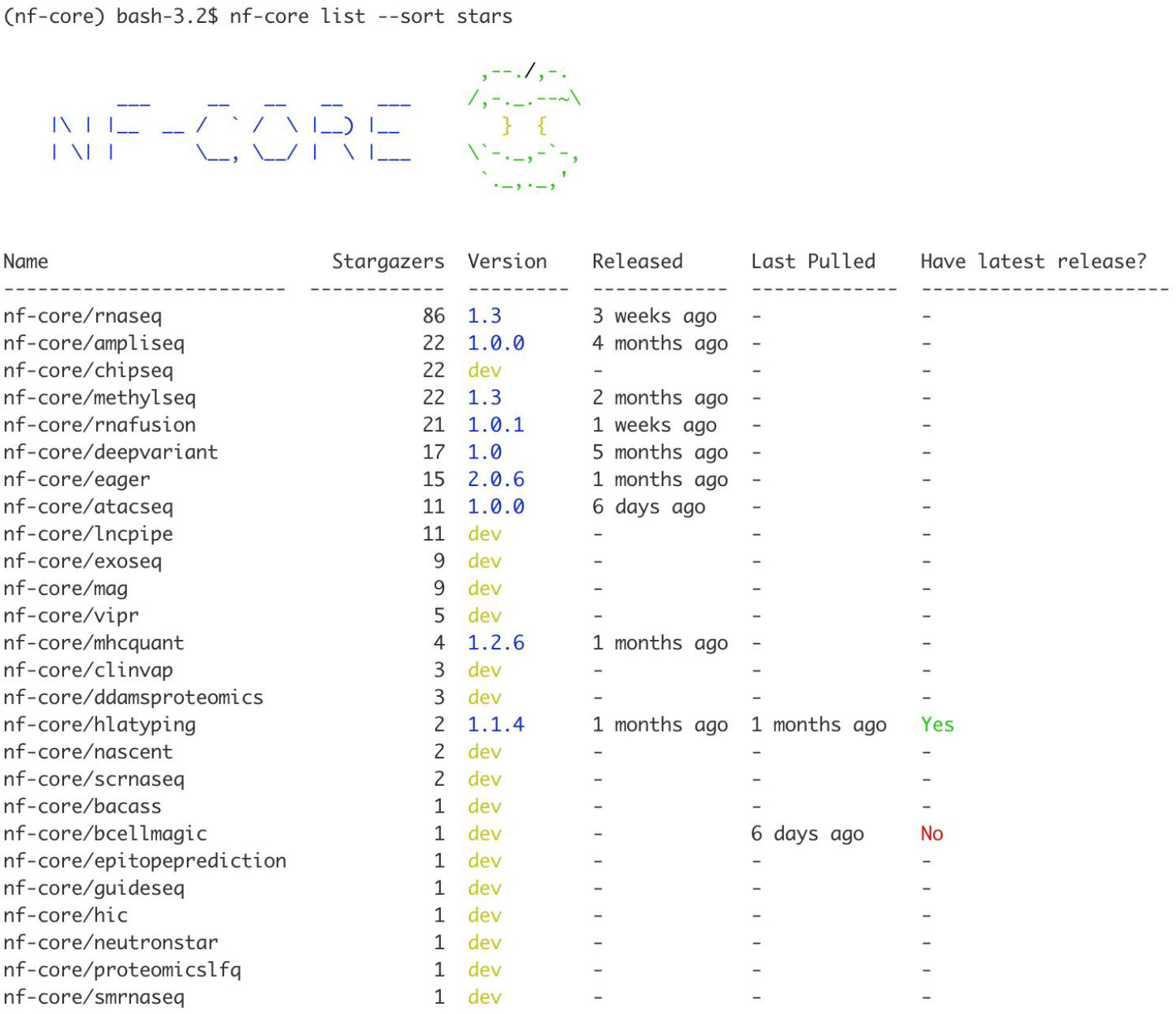
Example output from the nf-core framework tools, the ‘list’ command shows available analysis pipelines.

**Table 1.**
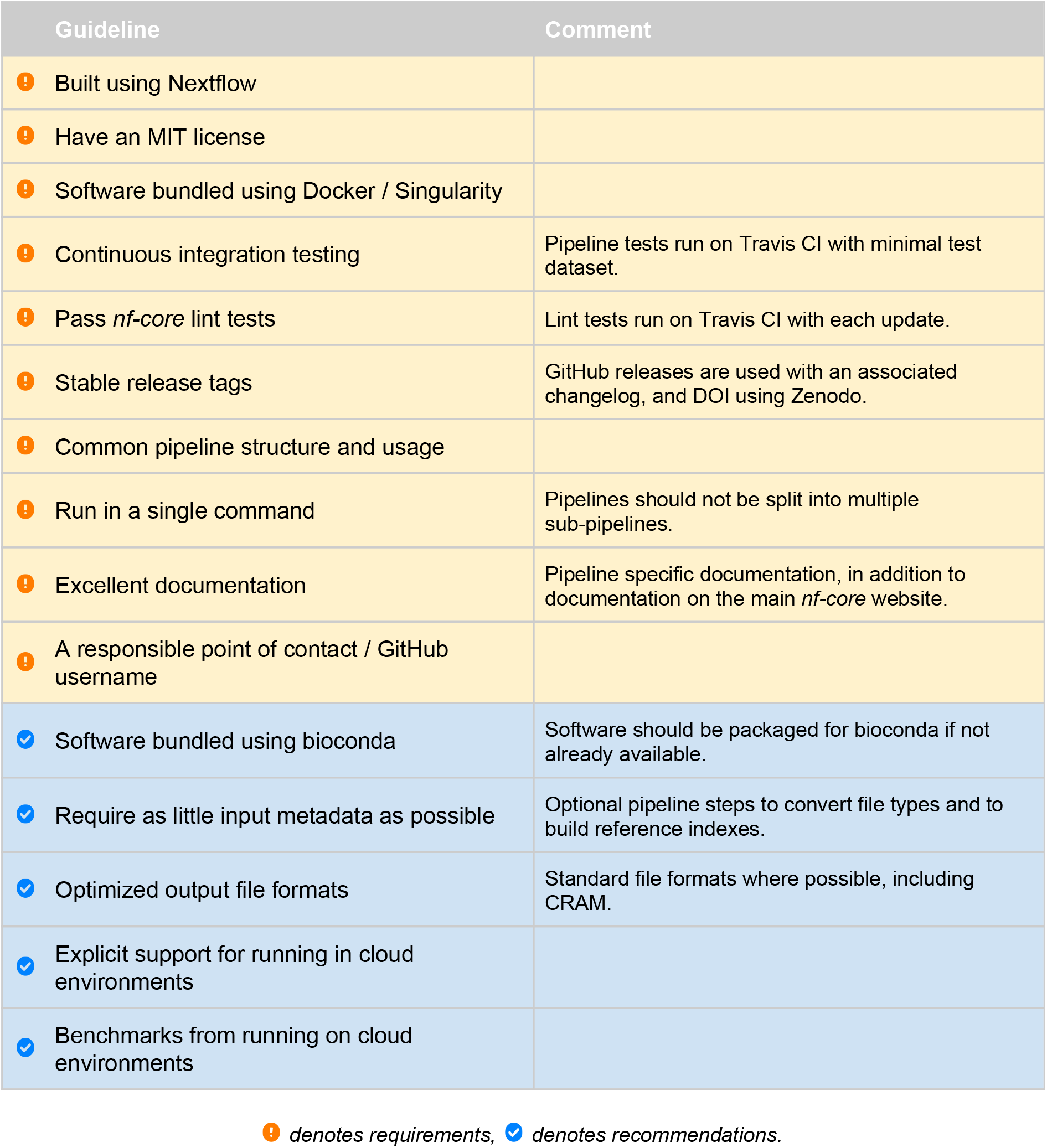
nf-core guidelines. See https://nf-co.re/developers/guidelines for the latest version.

To help developers get started with new pipelines, *nf-core* provides a standardized template pipeline which adheres to all *nf-core* guidelines (Table 1). The use of this template greatly lowers the learning curve when getting started with Nextflow and helps to homogenize the coding style of all *nf-core* pipelines.

The *nf-core* framework changes over time and new best practice insights need to be integrated into existing pipelines. We provide an automated synchronization mechanism that distributes relevant *nf-core* template changes to existing pipelines at every framework release (Fig S1). The pipeline maintainers can review the suggested changes and merge them into the development version of the source code, updating the pipeline with minimal manual effort.

All *nf-core* pipelines can be run on any computing infrastructure supported by Nextflow. This facilitates the possibility to run *nf-core* pipelines on most on-premise as well as cloud computing infrastructures such as AWS Batch, Google Cloud Platform and Kubernetes. The opportunity to run pipelines locally during development for initial examination, and then proceed seamlessly to large-scale computational resources in HPC or cloud settings provides users and developers with great flexibility.

In summary, the utilization of workflow management systems and container technologies alone do not ensure reproducible computational analysis. They can still lead to local platform dependencies which can inherently limit portability and reproducibility. *nf-core* provides both an architectural framework and a community platform to develop, share and discuss best-practice bioinformatics analysis pipelines. By leveraging the full feature sets of Nextflow, Conda, Docker, and Singularity, the *nf-core* framework directly tackles the reproducibility and portability concerns of researchers. This helps new users and developers alike, and enables the sharing of best-practice implementations across institutions. All pipelines adhere to the same usage pattern enabling both developers and users to familiarize themselves with *nf-core* pipelines with minimal effort.

As the usage of workflow management tools spreads, an increasing number of tertiary tools are tying into the ecosystem. The *nf-core* analysis pipelines are at the forefront of this, collaborating with initiatives such as bio.tools^28^, the GA4GH-compliant Dockstore^29^, and with plans to work together with the Biocontainers^18^ project in the future to further simplify software packaging.

We are in the process of building an interactive command-line and graphical user interface to further simplify the launching of *nf-core* pipelines. Other plans include extending the existing framework for automated benchmarking using full-size test datasets, the development of tools for the provision of more accurate cloud computing price estimates, and the implementation of the native Nextflow modular language across *nf-core* pipelines. Looking ahead, we hope to welcome more contributors and pipelines to the *nf-core* community to build on the solid foundation that has already been established.

## Acknowledgments

The authors would also like to thank the following people for their contributions: C Wang, S Paneerselvam, F Bonath, O Contreras-López, S Haglund, R Hammarén, D Yuen, J Wan, M Proks, A Jemt, RA Olsen, J Westholm (Science for Life Laboratory, Stockholm, Sweden); Chi Chuan (A*STAR Genome Institute of Singapore, Singapore, Singapore); M Hoeppner (Institut für Klinische Molekularbiologie, Kiel, Germany); V Malladi (University of Texas Southwestern Medical Center, Dallas, USA); A Duncan (Ontario Institute for Cancer Research, Canada); H Gourlé (Swedish University of Agricultural Sciences, Uppsala, Sweden); S Heumos, C Mohr, D Straub, T Koch, G Gabernet (Quantitative Biology Centre, Tübingen, Germany); G Kelly (Francis Crick Institute London, UK); Q Zhao (Sun Yat-sen University Cancer Center, Guangzhou, China); and E Floden (Centre for Genomic Regulation (CRG), Barcelona, Spain). AP, SF and SN acknowledge funding from Deutsche Forschungsgemeinschaft (core facilities initiative, KO-2313/6-1 and KO-2313-2, Institutional Strategy of the University of Tübingen, ZUK 63). AP and SN acknowledge funding by the Sonderforschungsbereich SFB/TR 209 “Liver cancer” of the Deutsche Forschungsgemeinschaft (DFG).

## Supplementary Materials

**Figure S1:**
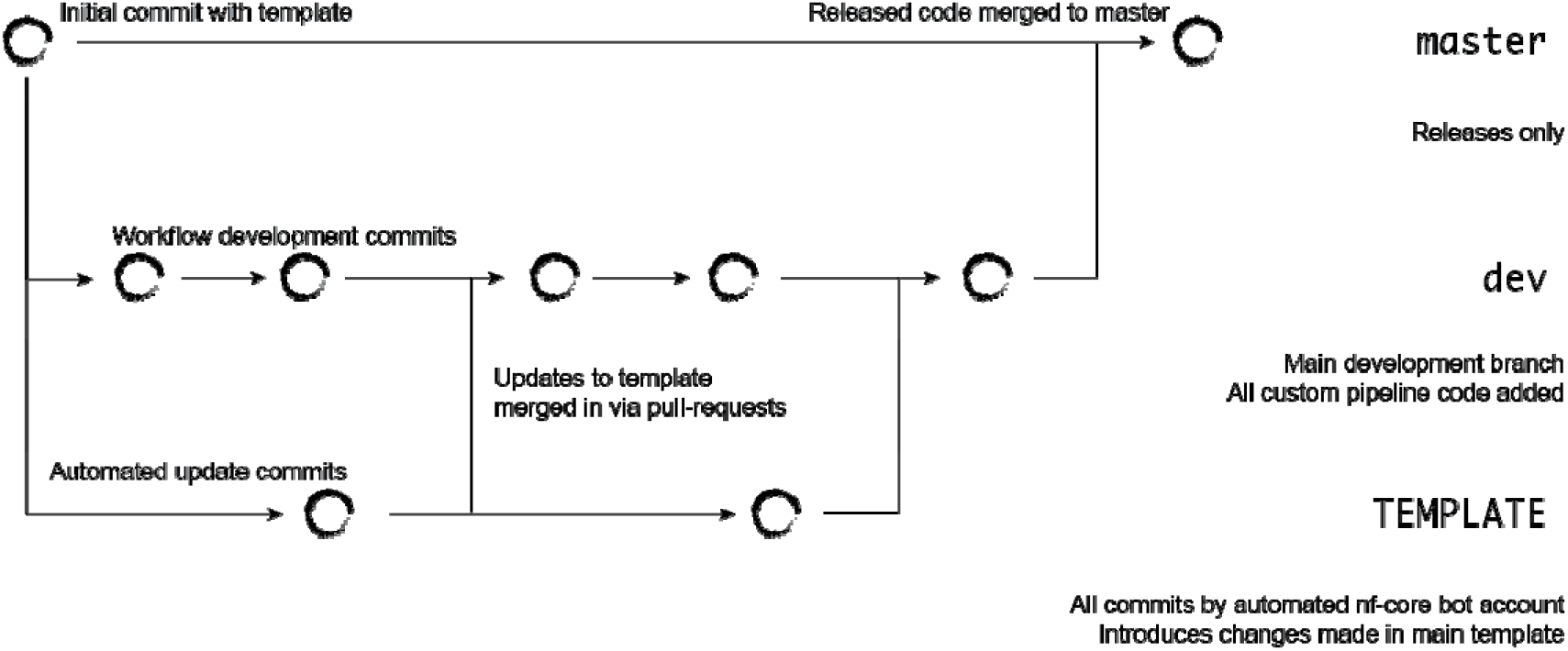
Template synchronization works using Git branching: the first commit for a pipeline is created immediately after using the template creation command and is branched into a special `TEMPLATE` branch. This remains untouched by the developer to keep the template commit history clean. Every time a new helper tool version gets released the continuous integration script automatically creates the pipeline again using the new version of the template. This is committed to the `template` branch and a pull-request to the pipeline’s development branch is automatically created with a Github bot account. This pull-request will only contain the relevant diff produced by the updated template. By merging the pull-request the changes in the pipeline’s implementation remain untouched but framework changes are integrated.

## Notes

#### Summary of Updates

Fixed PDF output, should be without warped text now.

